# Relating growth potential and biofilm formation of Shigatoxigenic *Escherichia coli* to *in planta* colonisation and the metabolome of ready-to-eat crops

**DOI:** 10.1101/523175

**Authors:** Bernhard Merget, Ken J. Forbes, Fiona Brennan, Sean McAteer, Tom Shepherd, Norval J.C. Strachan, Nicola J. Holden

**Affiliations:** Cell and Molecular Sciences, The James Hutton Institute, Dundee, DD2 5DA, UK; School of Biological Sciences, The University of Aberdeen, Cruickshank Building. St Machar Drive, Aberdeen, AB24 3UU, UK; School of Medicine and Dentistry, The University of Aberdeen, Foresterhill, Aberdeen, AB25 2ZD, UK; Teagasc, Dept. of Environment, Soils and Land-Use, Johnstown Castle, Wexford, Republic of Ireland; Roslin Institute & R(D)SVS, The University of Edinburgh, Easter Bush, EH25 9RG, UK

**Keywords:** spinach, lettuce, alfalfa, fenugreek, *E. coli* O157:H7, EHEC

## Abstract

Contamination of fresh produce with pathogenic *Escherichia coli,* including Shigatoxigenic *E. coli* (STEC), represents a serious risk to human health. Colonisation is governed by multiple bacterial and plant factors that can impact on the probability and suitability of bacterial growth. Thus, we aimed to determine whether the growth potential of STEC for plants associated with foodborne outbreaks (two leafy vegetables and two sprouted seed species), is predictive for colonisation of living plants as assessed from growth kinetics and biofilm formation in plant extracts. Fitness of STEC was compared to environmental *E. coli*, at temperatures relevant to plant growth. Growth kinetics in plant extracts varied in a plant-dependent and isolate-dependent manner for all isolates, with spinach leaf lysates supporting the fastest rates of growth. Spinach extracts also supported the highest levels of biofilm formation. Saccharides were identified as the major driver of bacterial growth, although no single metabolite could be correlated with growth kinetics. The highest level of *in planta* colonisation occurred on alfalfa sprouts, though internalisation was 10-times more prevalent in the leafy vegetables than in sprouted seeds. Marked differences in *in planta* growth meant that growth potential could only be inferred for STEC for sprouted seeds. In contrast, biofilm formation in extracts related to spinach colonisation. Overall, the capacity of *E. coli* to colonise, grow and internalise within plants or plant-derived matrices were influenced by the isolate type, plant species, plant tissue type and temperature, complicating any straight-forward relationship between *in vitro* and *in planta* behaviours.

**Importance:** Fresh produce is an important vehicle for STEC transmission and experimental evidence shows that STEC can colonise plants as secondary hosts, but differences in the capacity to colonise occur between different plant species and tissues. Therefore, an understanding of the impact of these plant factors have on the ability of STEC to grow and establish is required for food safety considerations and risk assessment. Here, we determined whether growth and the ability of STEC to form biofilms in plants extracts could be related to specific plant metabolites or could predict the ability of the bacteria to colonise living plants. Growth rates for sprouted seeds (alfalfa and fenugreek) exhibited a positive relationship between plant extracts and living plants, but not for leafy vegetables (lettuce and spinach). Therefore, the detailed variations at the level of the bacterial isolate, plant species and tissue type all need to be considered in risk assessment.

## Introduction

Contamination of fresh produce from Shigatoxigenic *Escherichia coli* (STEC) presents a serious hazard as a cause of food-borne illnesses, diarrhoea and enterohemorrhagic disease. Fresh produce is a major vehicle of transmission of STEC, with foods of plant origin accounting for the majority of *E. coli* and *Shigella* outbreaks in the USA (50). Fresh produce is often eaten raw or minimally processed and contamination of the produce can occur at any point along the food chain from farm to fork, with major outbreaks e.g. spinach (30) and sprouted seeds (6). STEC has been shown to interact with plants and can use them as secondary hosts (16, 25), which has implications for pre-harvest contamination, as well as persisting on post-harvest produce (29, 31, 32, 34, 52).

Colonisation of host plants by *E. coli* is governed by a range of environmental, bacterial and plant factors. Initial contact and attachment of bacteria on plant tissue is defined by motility, adherence factors and plant cell wall components (33, 64, 65), while establishment is influenced by a range of plant biotic (44, 67, 68) and abiotic factors (15, 62). The ability of bacteria to grow in the presence of plant material is a key factor in assessing risk, and although proliferation is well known to be influenced by physio-chemico factors (5, 23, 57, 58), risk assessments for STEC on fresh produce tend to consider plants as a homogenous whole (12, 21, 51).

STEC preferentially colonise the roots and rhizosphere of fresh produce plants over leafy tissue and have been shown to internalise into plant tissue, where they can persist in the apoplastic space as endophytes (13, 72). The apoplast contains metabolites, such as solutes, sugars, proteins and cell wall components (54) and as such provides a rich environment for many bacterial species, both commensal bacteria and human pathogens (20, 28). The rate of STEC internalisation is dependent on multiple factors including the plant species and tissue (73) and how plants are propagated (17-19). Specificity in the response of STEC to different plant species and tissue types has been demonstrated at the transcriptional level (9, 38). Therefore, there is a need to take into account specificity of the STEC-plant interactions that could impact risk.

Determination of the growth potential of a bacterial population takes into account the probability of growth together with the suitability of the growing population for a particular environment (22). It is used as a measure in risk assessment, e.g. for growth of STEC in water (70). In plant hosts, bacterial growth potential is governed by several factors, including bacterial growth rates, initial adherence and colony establishment, which is often in biofilms, as well as plant-dependent factors including metabolite availability and the ability to withstand plant defence responses (26). Therefore, the aim here was to determine if *in vitro* growth kinetics and biofilm formation of STEC in plant extracts, together with plant metabolite analysis, could be related to colonisation of plants that are associated with food-borne outbreaks, and hence inform on growth potential of STEC *in planta*. Use of genetically distinct *E. coli* isolates (two STEC, two environmental and one laboratory isolate) enabled assessment of phenotypic variation within plants or plant-derived matrices to be compared. Growth kinetics and biofilm formation were quantified in different tissue extracts of two leafy vegetables, lettuce and spinach, and two sprouted seeds, fenugreek and alfalfa sprouts. Growth kinetics was related to metabolomics of the extracts. Quantification of *in planta* colonisation and internalisation allowed a correlation analysis the for two STEC isolates.

## Results

### Growth rate parameterisation

To relate growth potential to colonisation of STEC in fresh produce plants, *in vitro* growth rates were first measured in plant extracts. Representative edible species associated with food-borne outbreaks were used: two leafy greens (lettuce, spinach) and two sprouted seeds (fenugreek, alfalfa). Plant tissues used were to represent edible, non-edible and internalised tissues of the leafy greens from total lysates of leaves or roots, and apoplastic washing recovered from leaves, respectively, while total sprout lysates were used to represent edible sprouts. A panel of five *E. coli* was assessed (Table 1) to allow relative fitness of two STEC O157:H7 Stx-isolates derived from fresh produce-associated outbreaks to be compared to two environmental isolates from plant roots and soil. A K-12 faecal-derived and laboratory-adapted isolate was included for reference. Growth was assessed at three temperatures (18, 20 and 25°C) to represent relevant growth temperatures of leafy greens and sprouted seeds. Growth kinetics were measured from optical densities derived from a plate reader (as described by others (22).

**Table 1.** Bacterial isolates used in this study ST = sequence type, Stx = Shiga toxin presence, nd = not determined, n/a = not applicable. Isolate Sakai used here is the *stx*-inactivated derivative (10). * Isolate ZAP1589, derived from H110320350 (Perry et al., 2013) has both *stx*-encoding regions removed, and is H7 positive but non-motile.

| Isolate Name | Serotype | ST | Stx | Source | Reference |
| --- | --- | --- | --- | --- | --- |
| MG1655 | OR:H48 | 98 | n/a | faecal/lab | (24) |
| JHI5025 | nd | 2055 | n/a | soil | (27) |
| JHI5039 | nd | 2303 | n/a | root | (27) |
| Sakai | O157:H7 | 11 | negative | sprout / | (10) |
| ZAP1589 | O157:H7 | 11 | negative | leek / clinical | (53)* |

### *E. coli* growth rates in plant extracts

Primary modelling of *in vitro* growth data in plant extracts successfully fitted 86.7 % (117 of 135) growth curves with a non-linear Baranyi model (SM1). Mis-fits were improved by manually truncating the growth curves to before the observed decrease in cell density that occurred in stationary phase, resulting in R^2^_adj_ = 0.996 (Fig. S1, Table S1). Comparison of the maximum growth rates (µ) showed highest growth rates in spinach extracts, with fastest growth in leaf lysates at 18 ° C or apoplast at 25 °C (Fig. 1A), while in lettuce the fastest growth occurred in apoplastic extract at all temperatures tested (Fig. 1B). All isolates grew consistently faster in fenugreek sprout extracts than in alfalfa, and either sprout extract supported faster growth than defined medium (RDMG) (Fig. 1C). The *E. coli* O157:H7 isolates showed differential responses in the different extracts and their growth rates were as fast or faster than the environmental isolates in almost all extracts. The lowest growth rates occurred for the laboratory-adapted isolate MG1655. The plant extract tissue-type as well as the bacterial isolate significantly impacted μ, from a two-way ANOVA at 18 °C (F (4, 7363) = 76.3; p < 0.0001 and F (8, 7363) = 436.4; p < 0.0001, for bacterial isolate and extract type, respectively) and at 20 °C (F (4, 8387) = 160.3; p < 0.0001 and F (8, 8387) = 416.1; p < 0.0001, for bacterial isolate and extract type, respectively).

**Figure 1.**
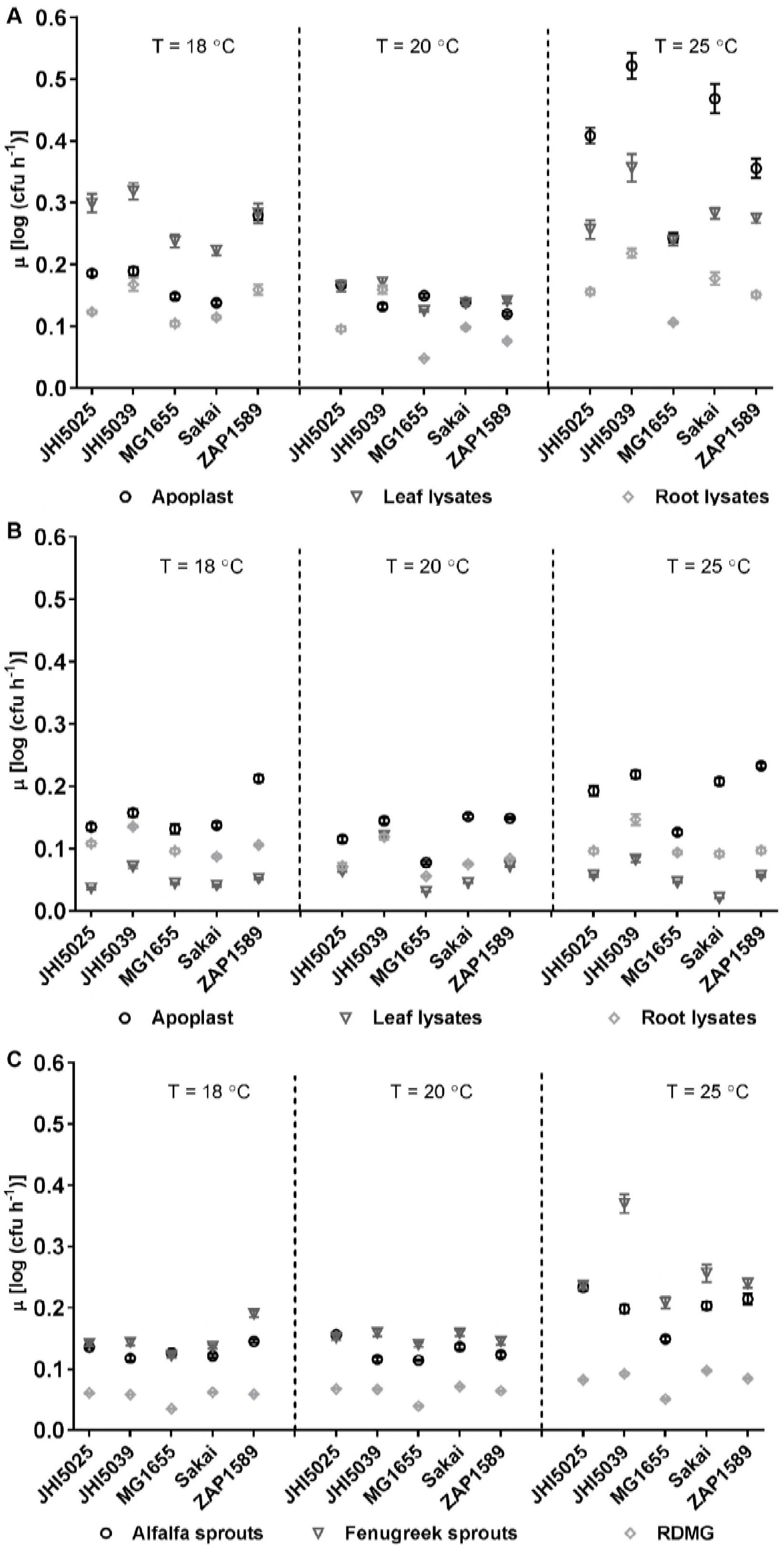
Maximum growth rates (μ) of reference *E. coli* isolates in plant extracts. Maximum growth rates (μ) were calculated using the Baranyi model for the reference *E. coli* isolates in spinach **(A)** or lettuce **(B)** aploplast (circles), leaf lysates (triangles) and root lysates (diamonds) extracts, or in alfalfa (circles) or fenugreek (triangles) sprouts lysate extracts **(C)** with RDMG (diamonds) as no-plant extract control, at 18, 20 or 25 °C. Each point is the average rate (n = 12), with standard errors indicated by bars.

Growth was almost always highest at 25 °C, although with exceptions, e.g. for *E. coli* O157:H7 isolate ZAP1589 in lettuce extracts. Growth characteristics were similar at both 18 and 20 °C, but μ were in general lower at 20 °C than at 18 °C. This counterintuitive result was reproducible and occurred in all growth experiments. It meant that secondary modelling for temperature was not possible. It was possible, however, for temperature-effects of growth in the defined medium without plant extracts, which produced a linear distribution for temperature for all five *E. coli* isolates (R^2^ = 0.996 to 1) (SM2), indicating the effect was due to the plant extracts and not a systemic error.

### Metabolite analysis of fresh produce plant extracts

To establish the impacts of different plant components on the growth of the *E. coli* isolates, metabolite analysis was determined for the extracts. Detection of absolute levels of mono-and disaccharides (sucrose, fructose, glucose, arabinose) showed the highest abundance in fenugreek sprout extracts, followed by lettuce apoplast and lettuce leaf lysates (Table 2). Sucrose was the most abundant sugar in all species and cultivars, except for alfalfa, which had high levels of fructose and glucose. Arabinose was only detected in the apolastic fluid of spinach and lettuce, accounting for 0.36 % and 0.23 % of all sugars, respectively. A two-way ANOVA found significant differences for tissue types (F (7, 60) = 16.5; p < 0.0001).

**Table 2.**
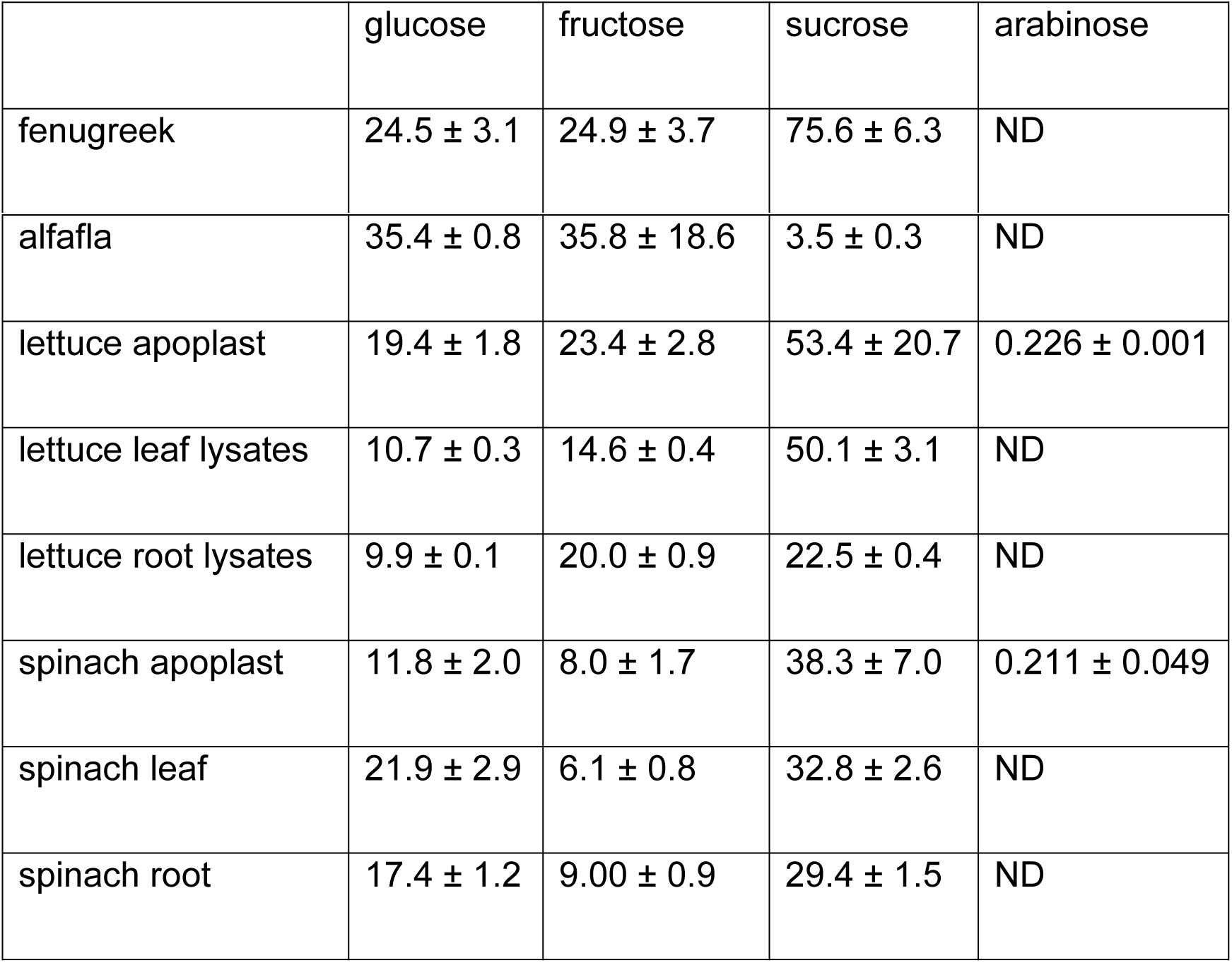
Quantification of saccharides from plant extracts Concentrations of mono-and disaccharides determined by HPLC (μg mg^−1^). ND – not detected.

The levels of amino acids and other metabolites were determined from identification of 116 polar metabolites, of which 60 were assigned and mapped onto a simplified polar metabolite pathway for plants to visualise metabolite availability for the bacteria (Fig. S2). The abundance ratio of each compound against the internal standard ribitol, generated a response ratio (RR) to allow semi-quantitative comparison (Table S2). Differences occurred between species and tissue types in a similar pattern to the mono-and disaccharides (Table 2), and for 12 metabolites including fructose, glucose and sucrose, there were significantly different RR (two-way ANOVA and Tukey multiple comparison, F (7, 854) = 37.2, p < 0.0001). Small amounts of arabinose could be found in all tissues with no significant differences between host species or tissue types. Grouping metabolites by structure (Fig. 2A) for monosaccharides, polysaccharides, amino acids, organic acids and other metabolites, showed that the highest total saccharides were present in fenugreek sprouts, while alfalfa was higher in monosaccharides and amino acids. The organic acids in spinach apoplast consisted mainly of oxalic acid, which was almost double the amount in spinach leaf lysates. The percentage composition showed that the majority of metabolites in all lettuce extracts are polysaccharides, compared to mainly of organic acids in all spinach extracts.

**Figure 2.**
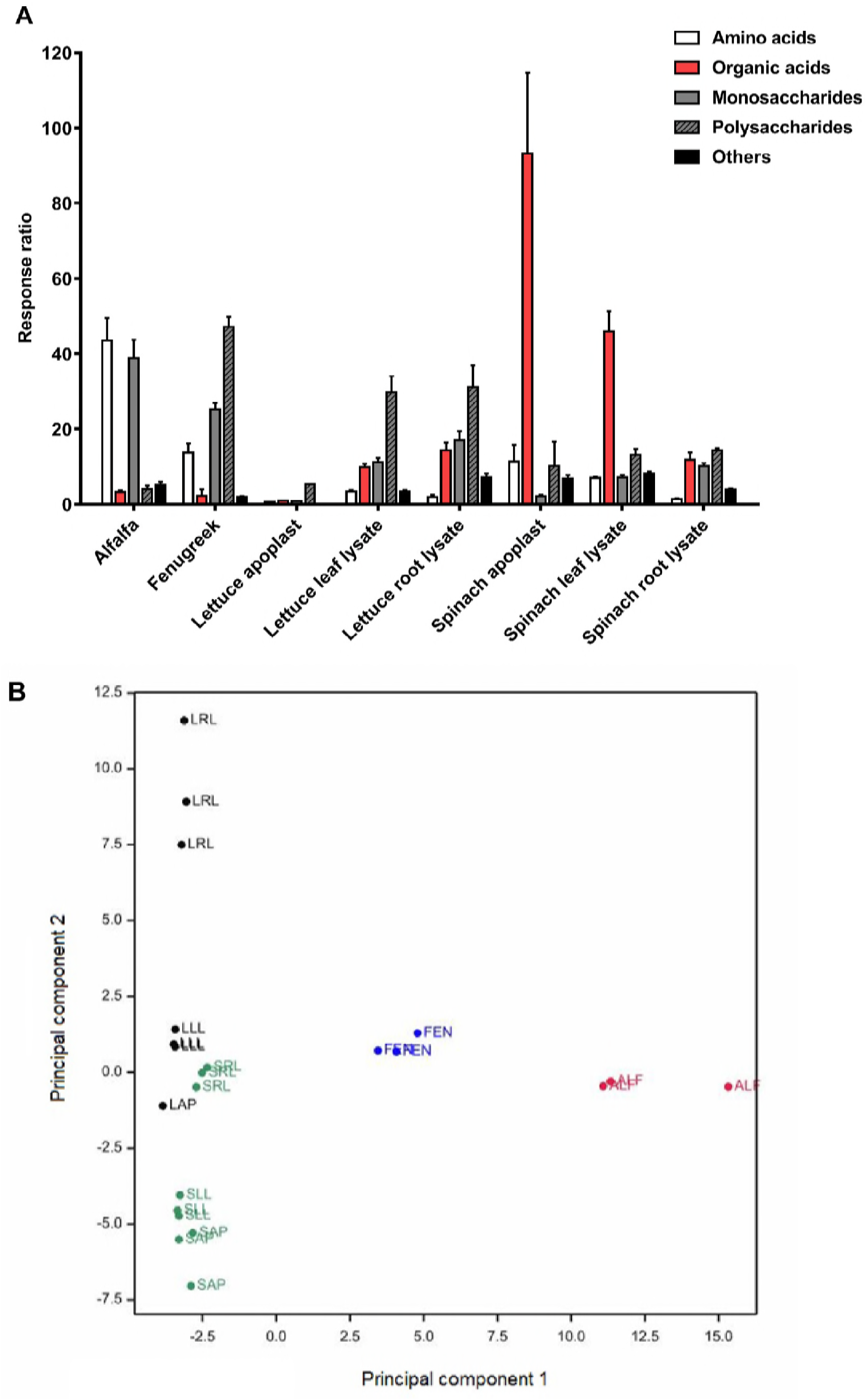
Plant extract metabolomics and grouping. The 60 assigned metabolites from all species and tissues are separated into amino acids, organic acids, mono-and polysaccharides and others **(A)** by their mean total response ratio (with SD indicated by bars). (**B)** Score plot of principal component 1 (31 % variance) and component 2 (19 %) for all 116 polar metabolites, for alfalfa (ALF) in red, fenugreek (FEN) in blue, spinach (SAP, SLL, SRL) green and lettuce (LAP, LLL, LRL) black.

Significant variation of the metabolite content occurred between plant tissues, as well as for and individual metabolites (two-way ANOVA from assuming a parametric distribution, F (420, 854) = 43.15; p < 0.001). A principal components analysis (PCA) showed that the first five components accounted for ~ 85 % of variance, and 50 % of the variance for all detectable polar metabolites (n=116) was attributed to PC1 and 2 (Fig. 2B). This was supported by significant positive correlation for leaf lysates and apoplast extracts of lettuce and spinach (R^2^> 0.97), a weak correlation for the root lysates based on species (R^2^ 0.542 – 0.757), with no significant correlation between any species for the tissues.

### The influence of plant extract metabolites on *E. coli* growth>

To relate any specific plant metabolites to bacterial growth, a correlation analysis was carried out between the plant extracts growth rates for two *E. coli* O157:H7 isolates (Sakai and ZAP1589) and the assigned metabolites. Several organic acids positively associated with maximal growth rates (µ), although there was a temperature-dependent effect. Metabolites associate with growth at 18 °C for isolate Sakai were galactosyl glycerol, threonic acid, and oxoproline (p ~ 0.04); at 20 °C, malic acid, fumaric acid and quinic acid (p = 0.014 – 0.048); and at 25 °C oxalic acid (p = 0.009), aspartic acid (p = 0.038), glutamic acid (p = 0.046), coumaric acid (p = 0.011) and uridine (p = 0.011). Chlorogenic acid (*trans*-5-O-caffeoyl-D-quinate) was consistently associated with growth for all temperatures (p 0.04 at 18 °C, p 0.004 at 20 °C, and p 0.04 at 25 °C). *E. coli* isolate ZAP1589 gave similar results, although there was also a bacterial isolate effect as there were no significant associations at 20 °C. Therefore, no single metabolite was identified as the major factor influencing *E. coli* growth rate, with a significant impact from growth temperature.

The main metabolite groups were then investigated as groups that could influence bacterial growth, by generating defined ‘artificial’ growth media comprising the main plant extract metabolites. The six most abundant metabolites were selected from lettuce apoplast or sprout extracts to represent contrasting metabolite profiles (Table 3). Each of the major groups of saccharides (SA), organic acids (OA) or amino acids (AA) were assessed independently by dilution, to restrict their effect, and at temperatures relevant to lettuce (18 °C) and sprouts (25°C). Maximal growth rates were similar in the sprout and lettuce extract artificial medium (Fig. 3), although reduced compared to the ‘complete’, natural extracts (Fig. 1). Growth rates were significantly reduced when the concentration of the saccharide group (SA) was reduced for both artificial media (all p < 0.0049), while restriction of the amino acids (AA) or organic acids (OA) had no impact (Fig. 3). The SA-dependent effect occurred for all *E. coli* isolates, although there were also significant isolate dependencies (two-way ANOVA, F (16, 28637) = 39.5; p < 0.0001 at 25 °C; two-way ANOVA, F (4, 9544) = 401.3; p < 0.0001 at 18 °C).

**Table 3.**
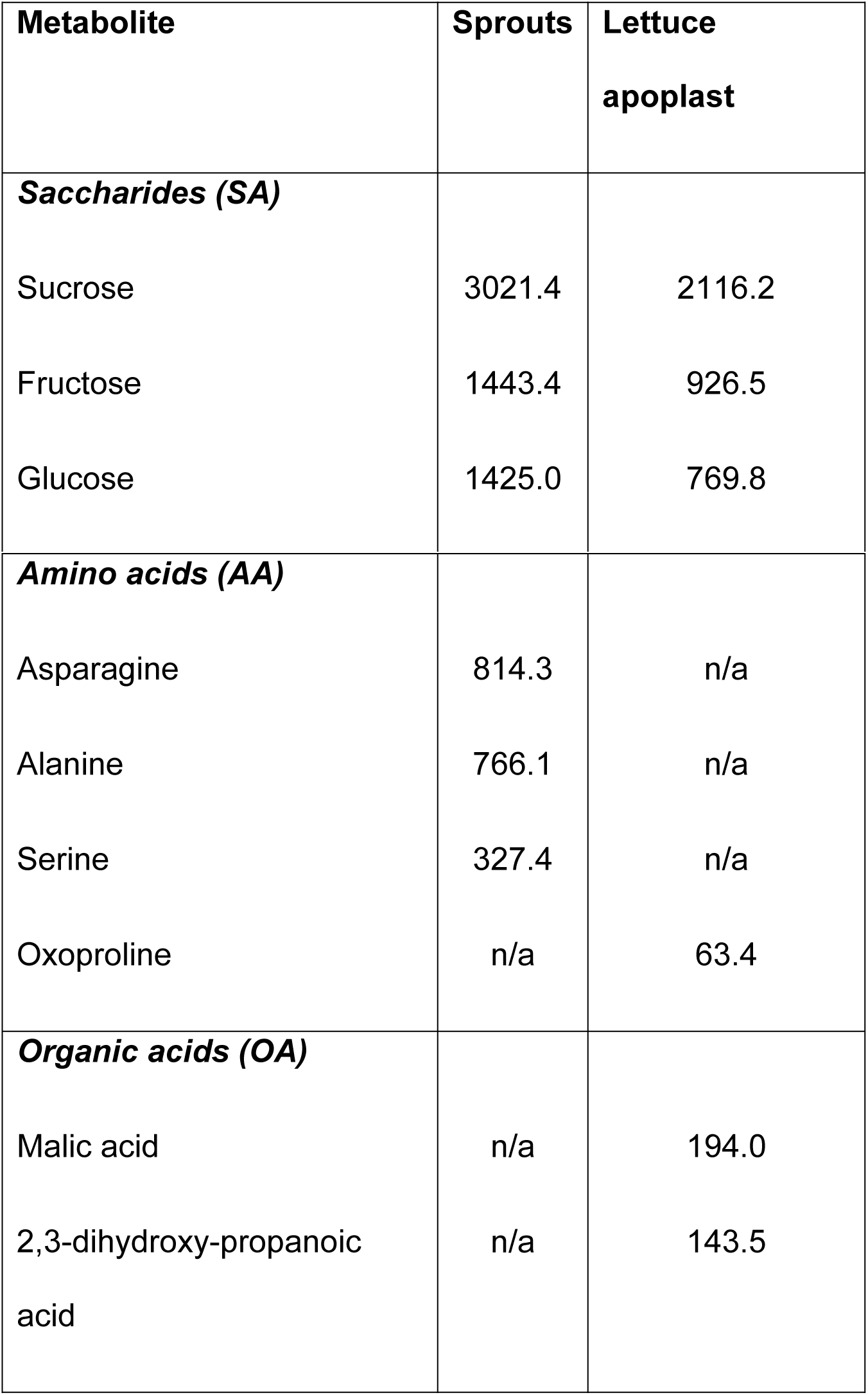
Composition of defined artificial media supplements Concentration (μg ml^−1^) as determined by HPLC and GC-MS for the major six components in sprout extracts (alfalfa and fenugreek combined), lettuce apoplast, used to generate defined ‘artificial’ media.

**Figure 3.**
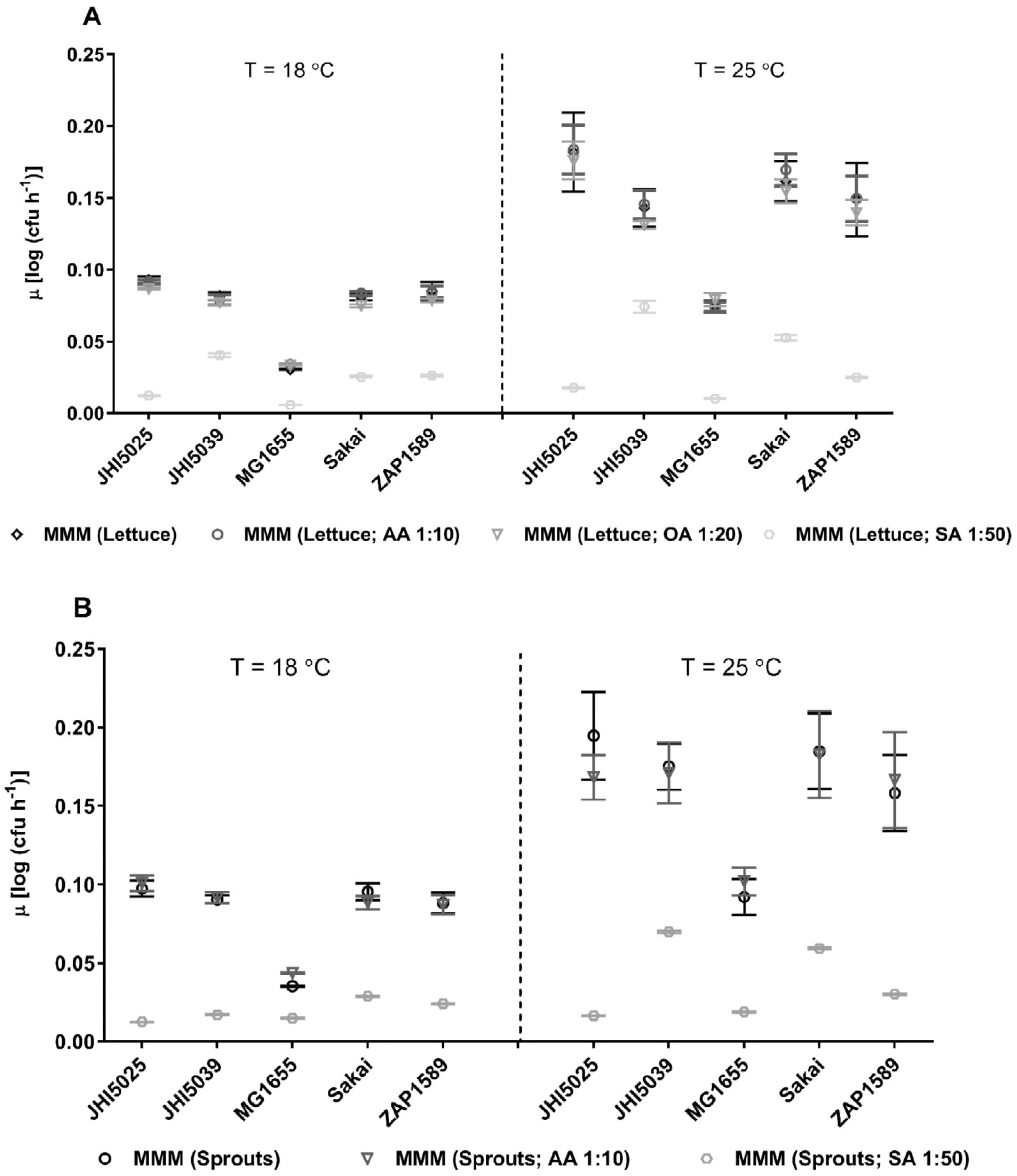
Maximum growth rates (μ) in artificial media mimicking plant extracts. Maximum growths rates (μ) calculated using the Baranyi model for the *E. coli* isolates at 18 C and 25 °C in media mimicking (**A**) lettuce apoplast or (**B**) sprout lysates (a mixture of alfalfa and fenugreek sprout metabolites) with specified dilutions. The base minimal MOPS medium (MMM) was supplemented with saccharides (SA), organic acids (OA) or amino acids (AA) at the dilution specified. Each point is the average rate with standard errors indicated by bars.

### The influence of plant extracts on *E. coli* biofilm formation

On host tissue *in planta*, bacterial colonies are more likely to be present in biofilms rather than as single cells. Therefore, the influence of the plant extracts of the leafy vegetables was tested for *E. coli* biofilm ability in isolation, i.e. on polystyrene surfaces. Spinach leaf lysates and root lysates were the only extracts that induced biofilm for all isolates, albeit minimal for isolate MG1655 (p < 0.0011, compared to isolate MG1655) (Table 4). The remaining extracts were not as conducive for biofilm formation, with the exception of one of the environmental isolates (JHI5025). This was not explained by different growth rates since this isolate did not exhibit the fastest growth rates in the extracts compared to the others (Fig. 1) and must therefore reflect increased adherence in the presence of the plant extracts. A qualitative risk ranking was determined for implementation of biofilm formation as a risk factor for the *E. coli* O157:H7 isolates (Sakai and ZAP1589) that identified spinach roots as the highest risk (from highest to lowest): spinach roots > spinach leaves > lettuce roots > lettuce leaves > spinach apoplast > lettuce apoplast.

**Table 4.**
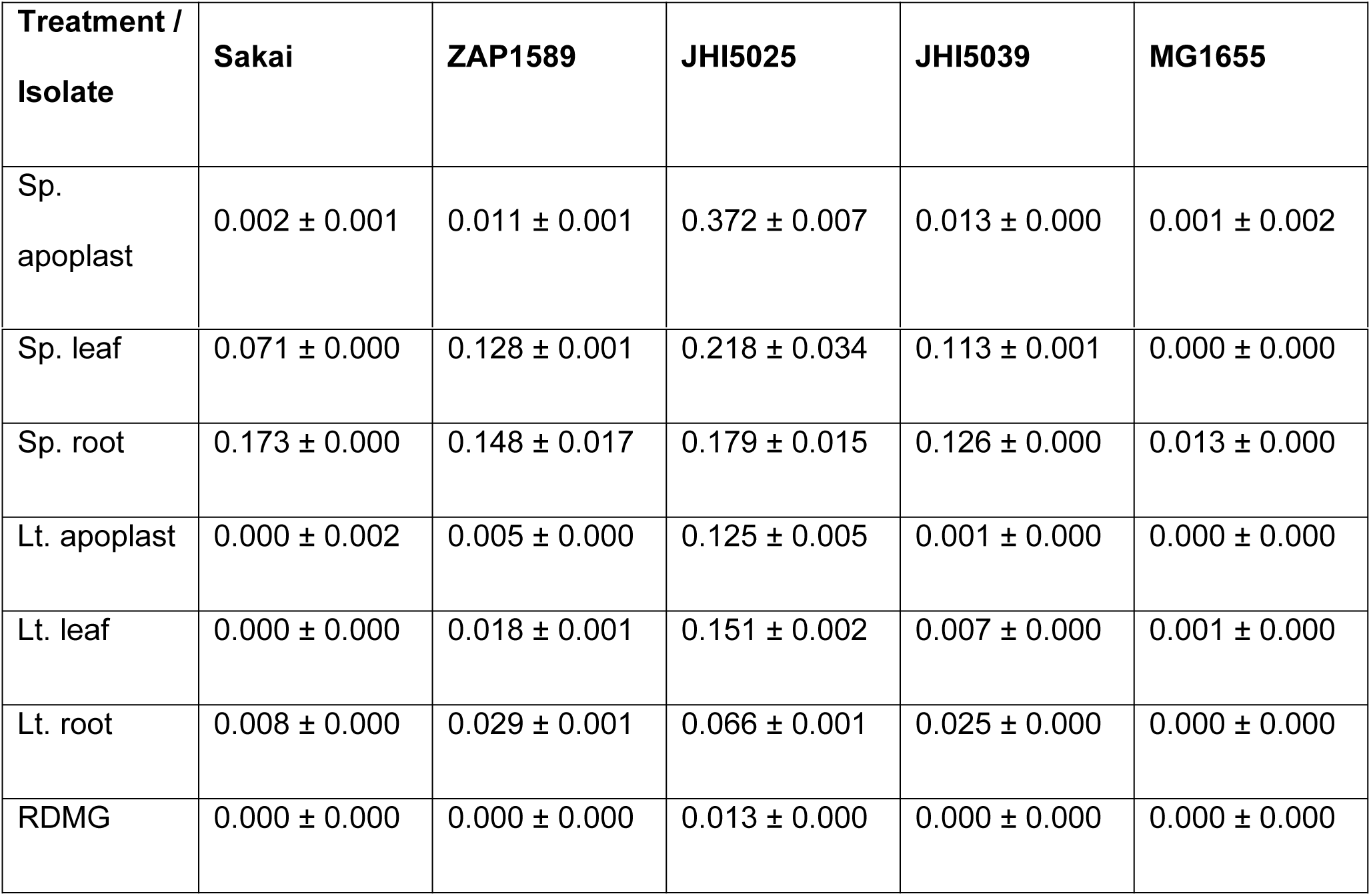
Biofilm formation for reference *E. coli* isolates in plant tissue extracts. Biofilms were formed on polystyrene multiwall plates following incubation in spinach (Sp.) and lettuce (Lt.) extracts (apoplast; leaf; root) and rich defined MOPS medium with glycerol (RDMG) at 18 °C, for 48 hrs in static conditions. The average (± variance) density of crystal violet at OD_590 nm_ is presented.

### *E. coli* O157:H7 colonisation and internalisation *in planta*

*E. coli* O157:H7 colonisation of leafy vegetables and sprouts was quantified to determine whether growth kinetics and biofilm formation in the extracts were predictive of *in planta* colonisation. Colonisation of the *E. coli* O157:H7 isolate (ZAP1589) was quantified on spinach and lettuce, and for both isolates (ZAP1589 and Sakai) on sprouted seeds. Our previous *in planta* data for lettuce and spinach plants showed that the highest levels of *E. coli* isolate Sakai occurred on spinach roots (73). Inoculation of spinach and lettuce with the high dose (10^7^ cfu ml^−1^) of *E. coli* isolate ZAP1589 also resulted in higher levels of bacteria on the roots compared to leaves, with similar levels on spinach and lettuce roots, e.g. 2.53 ± 0.97 and 2.69 ± 0.88 log (cfu g^−1^) at day 14, respectively (Fig. 4A, B). *In planta* colonisation of sprouted seeds by the two *E. coli* O157:H7 reference isolates was quantified for plants grown under conditions that mimic industry settings (hydroponics at 25 °C, three days) (Fig. 4C-F). A low inoculation dose of 10^3^ cfu ml^−1^ was used and total viable counts on day 0 were estimated by MPN since they fell below the direct plating detection threshold. Total counts of isolate Sakai increased by 4.5 log (cfu g^−1^) on alfalfa sprouts and 3 log (cfu g^−1^) on fenugreek sprouts, between 0 and 2 dpi. Viable counts for isolate ZAP1589 were generally lower on both sprouted seeds compared to isolate Sakai, but still reached 6.00 ± 0.253 log (cfu g^−1^) on alfalfa 2 dpi.

**Figure 4.**
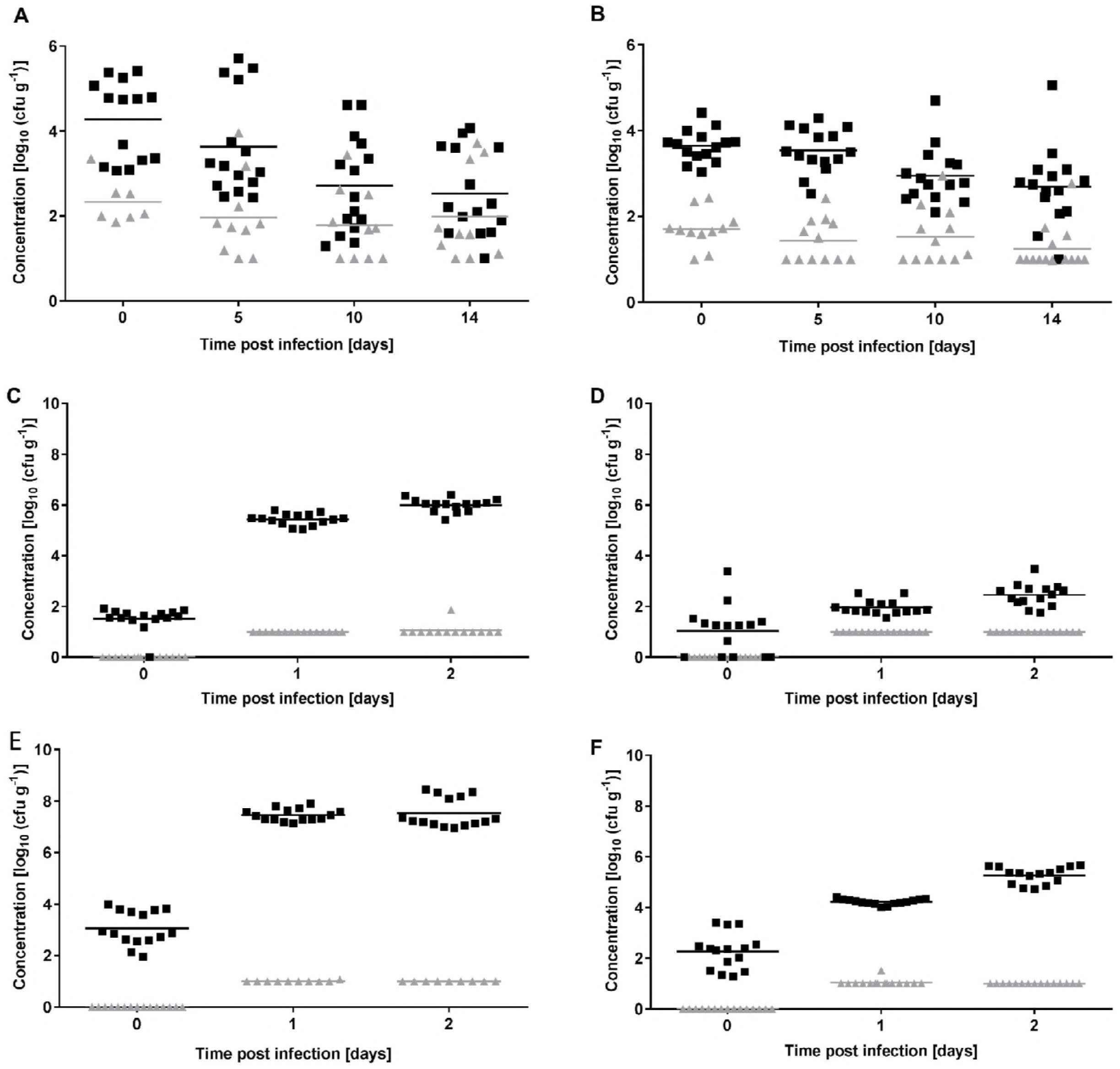
Total and internalised counts for *E. coli* O157:H7 *in planta*. The number of *E. coli* isolate ZAP1589 recovered from inoculation (10^7^ cfu ml^−1^) of (**A**) spinach (var. Amazon) or (**B**) lettuce (var. All Year Round) roots at 0, 5, 10 and 14 dpi.. The number of *coli* isolate ZAP1589 recovered from alfalfa (**C**) or fenugreek (**D**), and *E. coli* isolate Sakai recovered from alfalfa (**E**) or fenugreek sprouts (**F**), from inoculation at 10^3^ cfu ml^−1^, sampled at 0, 1 and 2 dpi.. Averages (lines) and individual samples counts are shown for the total (black) or internalised population (red) (n = 15: ~ 1.5 g per sample for sprouts, individual plants for spinach & lettuce). Sprout d0 data was assessed by MPN (level of detection = 0), otherwise minimum counts were manually levelled to the direct plating detection limit of 10 cfu g^−1^ on d1.

Internalisation was also assessed since endophytic behaviour is a feature of *E. coli* O157:H7 colonisation of fresh produce plants and growth potential could be reflected by growth in the apoplast washings. Internalisation of isolate ZAP1589 occurred to higher levels in spinach roots compared to lettuce roots (Fig. 4A, B), although the prevalence was similar in both plant species (60 % and 58.3 % of plants contained endophytic bacteria). In contrast, internalisation in sprouts only occurred on three occasions in all the experiments: isolate Sakai in alfalfa (1.07 log (cfu g^−1^)) and fenugreek (1.53 log (cfu g^−1^)) on day 1, and isolate ZAP1589 in alfalfa (1.87 log (cfu g^−1^)) on day 2. The prevalence was 7.1 % (1/14 samples positive), although the viable counts were close to the limit of detection by direct plating. Therefore, internalisation of *E. coli* O157:H7 isolates Sakai and ZAP1589 appeared to be a rare event on sprouted seeds, although they colonised the external sprout tissue to higher levels than on lettuce or spinach.

### Correlating *in planta* colonisation with plant extract growth rate kinetics

To relate growth kinetics in extracts with *in planta* growth, growth rates were estimated for *in planta* growth. This was possible for sprouted seeds since colonisation levels increased over time (Fig. 4). Alfalfa plants supported significantly faster growth rates for both *E. coli* O157:H7 isolates compared to fenugreek, at 2.23 ± 0.213 log cfu g^−1^ per day (R^2^ = 0.720) and 1.50 ± 0.0913 log cfu g^−1^ (R^2^ = 0.863) for Sakai on alfalfa and fenugreek sprouts, respectively, and for isolate ZAP1589, rates of 2.24 ± 0.159 log cfu g^−1^ (R^2^ = 0.822) and 0.710 ± 0.116 log cfu g^−1^ (R^2^= 0.464) per day on alfalfa and fenugreek sprouts, respectively. The difference in growth rate between the isolates on fenugreek sprouts was significant (p < 0.0001). Although *in planta* growth rates for *E. coli* isolates Sakai were estimated on spinach tissues (leaves, roots or internalised in leaf apoplast) or lettuce (leaves, roots) from low inoculation dose (10^3^ cfu ml^−1^) (73) these were non-significant since growth over the 10 day period was minimal or completely constrained, with a high degree of plant-to-plant variation. Growth rate estimates were not made when a high starting inoculum was used since the colonisation levels decreased over time (Fig. 4).

Comparison of the *in planta* and extract growth rate estimates were made for both *E. coli* O157:H7 isolates on sprouted seeds (at 25 °C) or in spinach and lettuce (at 18 °C) (Fig. 5). A positive correlation occurred for growth rate estimates in the sprouted seeds (R^2^ = 0.516), although this was not significant. Since *in planta* growth in spinach or lettuce tissues was minimal, there was no correlation with growth rates in corresponding extracts. Therefore, the restrictions in bacterial growth that occurred with living plants meant that growth rates in extracts could not be extrapolated to *in planta* growth potential for leafy vegetables, but did bear a positive relationship for sprouted seeds.

**Figure 5.**
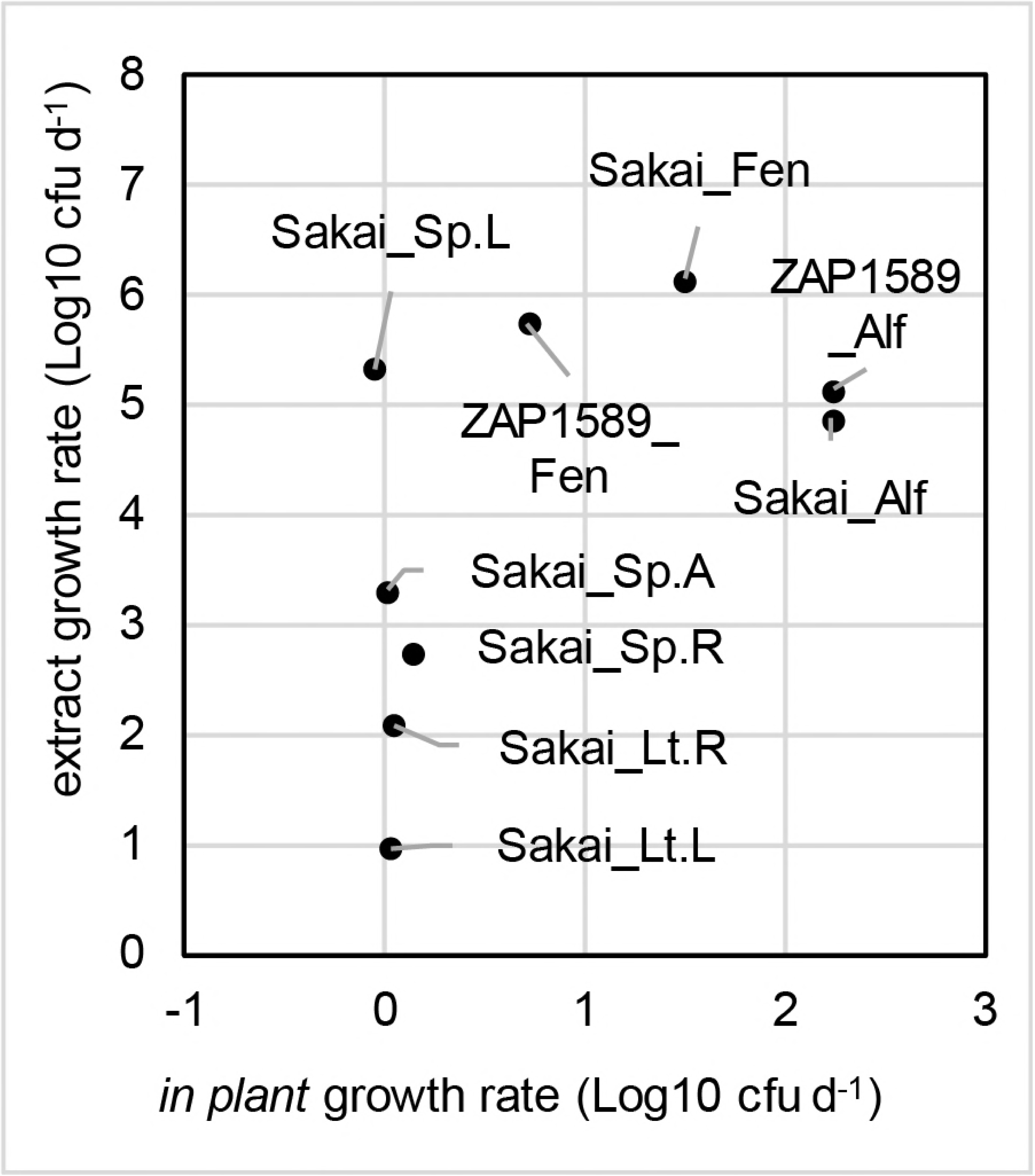
Comparison of *in planta* and extract growth rates for *E. coli* isolates Sakai and ZAP1589. Growth rates for *in planta* estimates were plotted against estimates for plant extract extracts, on a Log_10_ cfu day^−1^ basis for *E. coli* isolates Sakai and ZAP1589, normalised per g fresh weight for plant tissues or per ml for plant extracts. Estimates for sprouted seeds (alfalfa – Alf; fenugreek – Fen) were obtained for growth at 25 °C, and at 18 °C for spinach (Sp.) or lettuce (Lt.) tissues (apoplast – A; leaves – L; roots – R).

## Discussion

The potential for food-borne bacteria to grow in fresh produce food commodities is a key consideration in quantitative risk assessment. Factors that influence bacterial growth are the plant species and tissue, the bacterial species or isolate, and the surrounding environment. The growth potential of a bacterial population consists of proportion of the growing sub-population and the suitability of the environment for growth, and it provides a quantitative description of probability of growth (22). Therefore, the factors that influence growth potential of STEC in edible plants include plant-dependent and physio-chemico factors, as well as bacterial isolate-specific responses. Metabolically active components of plants can be extrapolated from plant extracts for bacterial growth dynamic measurements coupled with metabolite analysis. They also represent a bacterial growth substrate in their own right that could arise during the post-harvest production process e.g. from cut surfaces. A number of studies show growth of food-borne bacteria on plant extracts during the production process (35, 55, 56) and growth potential for *E. coli* O157:H7 has been evaluated in water (70). Here, maximum growth rates in plant extracts were strongly influenced by the plant tissue type and species, as well as the *E. coli* isolate tested and overlaid by temperature-dependent effects. *In planta* growth rates, however, was markedly different between the sprouted seeds and leafy vegetables, with a growth restriction evident in the leafy vegetables. The plant-dependent factors that could account for this difference include plant age, defence response, growth conditions and associated microbiomes. As such growth rates in the extracts could not be used to infer *in planta* growth potential for spinach or lettuce. In contrast, proliferation on sprouted seeds did bear a positive relationship to growth rates in extracts, although it was also dependent on the plant species and on bacterial isolate tested.

Saccharides were shown to be the major driving force for *E. coli* growth, which is unsurprising given their role in central metabolism (41). Although the levels of the most abundant sugars, glucose, fructose and sucrose (the disaccharide of glucose and fructose) could explain the high growth rates in sprout extracts, similarly rapid growth did not occur in lettuce leaf lysate extract, despite an abundance of sugars, indicating that plant species-specific inhibitory compounds exist. This is supported by the occurrence of more rapid growth rates in spinach leaf extracts compared to lettuce. Plant-dependent factors that could influence bacterial growth potential include the innate defence response (33), antimicrobial activity of plant secondary metabolites (71) and plant development stage (74).

Bacterial growth rates were not significantly impacted by manipulation of the major amino or organic acids from the extracts, although the phenolic acid, chlorogenate (*trans*-5-O-caffeoyl-D-quinate) was positively associated with growth. This contrasts to reports of its ability to inhibit fatty acid synthesis in *E. coli* isolate MG1655 (37) and prevent *E. coli* growth (75), but may be explained by differences in concentration between the extracts and exogenous application. Oxalate levels were relatively high in spinach, in keeping with previous reports that show an average as high as ~ 1000 mg / 100 g fresh weight (47) and correlated with growth for isolate Sakai at 25 °C. Amino acids levels were substantially higher in sprouted seed extracts compared to the leafy vegetables, which is likely a reflection of different developmental stages of the plants (4). It was notable that the artificial media did not support equivalent growth rates to the ‘complete’, natural extract media, indicating that other, minor nutrients in the extracts were utilised for maximal bacterial growth and also need to be accounted for in growth dynamics.

Bacteria including STEC, tend to form biofilms in association with plant tissue (11, 73, 74). Here, a risk ranking could be inferred from biofilm formation in the extracts, with spinach roots ranked highest. Curli is an important biofilm component for STEC associated with plants (7), but other biofilm components are likely to be responsible for the biofilm formation in extracts, since isolate Sakai did not form biofilms in spinach apoplast extract *in vitro* but does produce curli during endophytic colonisation and biofilm formation in leaves (73). This indicates that specific *in planta* cues induce different biofilm components. Alternative biofilm components that may be involved include Type 1 fimbriae, which was shown to be expressed by the environmental isolates JHI5025 and JIH5039 at 20 °C and promoted binding to spinach roots (40).

Internalisation of STEC into apoplastic spaces in plants presents a hazard as pathogens cannot be removed by conventional sanitation methods. However, growth potential for internalised *E. coli* O157:H7 could not be inferred from growth in apoplast extracts since endophytic proliferation was prevented or reduced in the apoplast (73). As the apoplast is a habitat for plant-associated endophytes (66) and phytopathogens (63), it appears that for *E. coli* additional factors such as the plant defence response need to be considered. The increased likelihood of internalisation into tissues of leafy vegetables compared to sprouted seeds for the *E. coli* O157:H7 isolates could be due to multiple factors including plant age, the competing microbiota and access to nutrients. Plant dependent factors have also been shown to impact colonisation of lettuce cultivars by STEC (Quilliam, et al. (59).

*In planta* colonisation of *E. coli* O157:H7 isolate Sakai was significantly higher than isolate ZAP1589, in both leafy tissue types and on both sprouted seed species (73). In contrast, growth rates in the plant extracts and in artificial media overlapped, albeit with specific extract-specific differences. Since isolate ZAP1589 was found to be flagellate but non-motile, this may reflect a role for flagella in plant colonisation (65). ZAP1589 growth rates on sprouted seeds were similar to the rates reported for other *E. coli* O157:H7 isolates on 2-day old alfalfa sprouts (8). Growth rates of both *E. coli* O157:H7 isolates in the extracts was, in general, as high as the environmental isolates, and almost always higher than the K-12 isolate, indicating similarities in fitness levels for STEC and environmental *E. coli*.

The ability of *E. coli* isolates to metabolise different carbon sources varies and could contribute to the isolate-dependent variations in growth rates. Although less than 50 % of *E. coli* isolates can metabolise sucrose (41), *E. coli* O157:H7 isolate Sakai encodes the sucrose transport genes (1) and sucrose degradation genes were expressed by this isolate on exposure to spinach extracts (9). The sucrose translocator from *S. enterica* serovar Typhimurium was expressed by a related epiphyte *in planta* (46). In contrast, fructose and glucose are sufficient sole carbon source-metabolites for *E. coli* and their role in bacterial metabolism is well characterised (41). An *E. coli* fructose metabolism gene has also been expressed in a related epiphyte *in planta* (36).

Growth rates normally positively correlate with temperature (60), as was observed for growth rates in the defined medium without plant extracts, which exhibited a linear distribution from 18 °C to 25 °C. However, maximal growth rates in the extracts were influenced in a non-linear manner by temperature. Similarly, a non-linear effect was reported in a meta-study on growth of STEC on lettuce (42). This could be due to *E. coli* adaption to the plant environment and resulting metabolic responses (9), and reflects the different organic acid correlations that occurred for different temperatures seen here. The implications are that a linear approximation, e.g. such as a Ratkowsky model, is not sufficient to describe *E. coli* growth in plant extracts, although it has been used to model growth on plants (43, 60, 61).

In conclusion, growth potential *in planta* was described in part, by growth rates in plant extracts, but only for sprouted seeds. On the other hand, biofilm formation in plant extracts showed some relation to *in planta* colonisation in leafy vegetables. Plant species-and tissue-type dependent differences in metabolites meant that no single metabolite could be correlated with growth, and the only positive association was with the combined group of saccharides. The marked differences in *in planta* colonisation between the sprouted seeds and leafy vegetables reinforces the higher risk associated with very young plants, grown under conditions conducive for bacterial growth (72). Therefore, although this data can inform hazard identification and risk analyses, it is evident that important specificities within each plant-microbe system need to be considered, and it is not possible to take a generalised view of STEC-plant colonisation.

## Materials and Methods

### Bacteria and media

The bacterial isolates panel comprised five isolates: two *E. coli* O157:H7 isolates, two environmental *E. coli* isolates and an *E. coli* K-12 isolate (Table 1). *E. coli* ZAP1589 is a Stx negative derivative, generated from isolate H110320350 (Methods SM3). Motility of isolate ZAP1589 and isolate H110320350 was tested on motility agar (0.7 %), and presence of the H7 flagella was confirmed by agglutination with the monoclonal H7 antibody.

Bacteria were cultured overnight in Lysogeny-broth medium (LB) at 37 °C (2, 3), with shaking at 200 rpm. Prior to experimentation an aliquot of the overnight culture was inoculated 1:100 in rich defined 3-(N-morpholino)propanesulfonic acid (MOPS) medium (48) with 0.2 % glycerol and essential and non-essential amino acids, termed ‘rich defined MOPS glycerol’ (RDMG), for 24 h at 18 °C and 200 rpm. Bacteria were collected by centrifugation, washed in phosphate buffered saline (PBS) and adjusted to the required starting optical density (OD) 600 nm. Media was supplemented with 30 μg ml^−1^ kanamycin, if required. Defined artificial ‘lettuce apoplast’ or ‘sprout extract’ media was generated by adding each group of constituents (Table 3) to a base minimal MOPs medium (MMM) lacking a carbon source and amino acids. Each component group was added at the defined concentration to represent the concentrations and composition present in lettuce apoplast or sprout extracts and by dilution of one major group at a time at: 1:50 saccharides (SA), 1:10 amino acids (AA) or 1:20 organic acids (OA), while the other groups were at 1:1. The pH of the sprout defined medium was 7.2 and lettuce apoplast defined medium 7.05. Viable counts were determined from 10-fold dilutions plated on MacConkey (MAC) agar, incubated overnight at 37 °C and counted manually the next day. All experiments were conducted in triplicate. Viable counts and OD_600_ nm were plotted in Excel 2010.

### Plant extracts and metabolite analysis

Lettuce (*Lactuca sativa*) var. All Year Round and spinach (*Spinacia oleracea*) var. Amazon were grown individually in 9 cm^3^ pots in compost for microbiological assays, or in vermiculite for metabolite analysis, in a glasshouse for three weeks. Fenugreek (*Trigonella foenum-graecum*) and alfalfa (*Medicago sativa*) seeds were soaked in sterile distilled water (SDW) for 3 h at room temperature (RT), surface sterilized with 3 % calcium hypochlorite (20,000 ppm ml^−1^ active chlorite) for 15 min, washed five times with SDW and soaked for 2 h in SDW at RT. Sprouts were transferred aseptically on distilled water agar (DWA) (0.5 % agar) and sprouted for two (alfalfa) or five (fenugreek) days at 25 °C in darkness. Leaf apoplastic washings were collected as described previously (Methods SM4), optimised for spinach and lettuce to minimize cytoplasmic contamination (39). All tissue extracts were made as described previously (9). In brief, vermiculite was gently washed off the roots with tap water and rinsed with SDW. Leaves and roots were separated with a sterile scalpel, macerated in liquid nitrogen with a pestle in a mortar and stored at −20 °C until use and pre-processed for sample clarification by mixing 1 g with 20 ml SDW, soaked on a shaker for 4 h, centrifuged at 5000 rcf for 15 min, and the supernatant heated to 50 °C for 30 min. The extract was centrifuged at 5000 rcf for 15 min and filter sterilised through a 0.45 μm filter for root tissue or 0.1 μm filter for leaf tissue. Sprouts were macerated in liquid nitrogen, processed as described above without a washing step to remove vermiculite, and filter sterilised through a 0.22 μm filter. Apoplast extracts were filtered sterilised through a 0.1 μm filter (Durapore, Merck, Germany). Extracts were made from ~ 5 plants per sample for leaves and roots and up to 24 plants for apolastic washings or for sprouts. 10 ml plant extract samples were used for GC-MS analysis as described in Methods SM5. Lysates were prepared for HPLC described previously by Shepherd, et al. (69).

### Bacterial growth rates

Bacterial growth rates were determined using a pre-warmed plate reader Bioscreen C plate reader (Oy Growth Curves Ab Ltd, Finland), set to different temperatures. The *E. coli* isolates were grown as described above, adjusted to an OD_600_ of 0.05 in PBS (~ 2.1 × 107 cfu ml^−1^) and inoculated at a 1:10 dilution in plant extracts (at 1:20 w/v in dH_2_O) or defined media (Table 3), in 200 μl total volume, in multi-well plates. Growth for the *E. coli* isolates was measured at 18, 20 and 25 °C in 100-microwell plates (Honeycomb, Thermo Fisher, USA). Wells were randomised in duplicate on the plate with negatives included. All growth curves in extracts were repeated three times with four replicates on plates. Measurements were recorded every 15 min for 48 hours and multi-well plates were shaken for 60 seconds pre-and post-measurement. Results were exported from plate reader proprietary software as tab-delimited files. For model fitting, 12 replicates of each isolate and medium type were averaged and converted to viable counts log (cfu h^−1^) (Methods SM6). A conversion factor of 4.2 × 10^8^ cfu ml^−1^ was applied so that all growth curves could be modelled using DM-Fit (Methods SM1). Secondary modelling was applied for different temperature as described (Methods SM2).

### Biofilms

Bacterial biofilms were measured as described previously by Merritt, et al. (45). Bacteria were grown aerobically in LB at 37 °C for 12 h, sub-cultured (1:1000 v/v) in RDMG for 18 h at 18 °C, diluted in PBS to OD_600_ of 0.05 and inoculated into plant extracts as per the growth rates determination in a 96 well polystyrene plate and incubated statically for 48 h at 18 °C. The washed wells were stained with 0.1 % crystal violet solution and solubilised with 95 % ethanol. The solution was transferred into a fresh plate and absorbance measured at 590 nm with a plate reader (Multiskan Go, Thermo Scientific, USA). Results were exported with the software SkanIt^TM^ (Thermo Scientific, USA) to Microsoft Excel 2010 for analysis.

### Plant colonisation assay

Lettuce and spinach plants (~ 3 weeks old) were transferred to a growth chamber (Snijders) at 21 °C; 75 % humidity and 16 h light – 8 h dark cycle (400 μE/m2.s (30.000 lux)) three days prior to inoculation and were not watered for ~ 18 h prior to inoculation. Roots were inoculated by placing pots in a plastic box containing a 1 litre suspension of *E. coli* Sakai or ZAP1589, diluted to OD_600_ of 0.02 (equivalent to 10^7^ cfu ml^−1^) in SDW, which partially submerged pots. After 1 h inoculation, the pots were transferred to the growth chamber until sampling. Sprouts were inoculated with 10^3^ cfu ml^−1^ bacteria in 0.5 l SDW for 1 h, rinsed with 0.5 × Murashige and Skoog (MS) basal medium (no sucrose), and transferred to petri dishes containing distilled water agar (DWA) (0.8 % agar) and incubated for up to three days at 25 °C. Negative controls were incubated with SDW without bacteria.

Lettuce and spinach roots were sampled at 0, 5, 10 and 14 days post infection (dpi), aseptically removed from aerial tissue with a sterile scalpel, the compost removed by washing with SDW, and the roots were transferred into 50 ml tubes, washed with PBS and the fresh weight determined. Sprouts were sampled at 0, 1, 2 dpi, where half were used to enumerate the total viable counts of *E. coli* and stored in PBS until further use (~ 30 min), and surface-associated bacteria were removed from the other half of the samples by surface sterilization with 200 ppm Ca(ClO)_2_ for lettuce/spinach roots or 20,000 ppm Ca(ClO)_2_ for sprouts, for 15 min. Surface decontamination of sprout tissue required at least 15,000 ppm of Ca(ClO)_2_ to eradicate external *E. coli*, but endophytes appeared to be protected from the active chlorite since endemic internalised bacteria occurred on recovery media after surface decontamination with 20,000 ppm Ca(ClO)_2_. The root/sprouts were washed five times with PBS to ensure removal of all loosely adherent bacterial cells and residual chlorine. Surface sterilisation was validated as described (73). Any samples containing surface-associated bacterial colonies were removed from subsequent analysis. Roots/sprouts were macerated using mortar and pestle in 2 ml PBS and ~ 50 mg sterile sand. The supernatant was diluted once for spinach and lettuce (1:1), three times for fenugreek (1:3) or four times for alfalfa (1:4) with PBS and 100 μl plated on MAC plates using a spiral plater (WASP, Don Whitley Scientific, UK) and incubated for 24 h at 37 °C. Plates were counted using a counting grid (WASP, Don Whitley Scientific, UK), multiplied by the dilution factor and converted to cfu ml^−1^. The experiment was repeated three times with five replicate samples per time point, and sprout samples comprised multiple (> 15) sprouts. The limit of detection from direct plating was 20 cfu ml^−1^, below which values were manually levelled to < 1 log (cfu ml^−1^) for lettuce and spinach root data. Since the level of inoculation of sprouts for day 0 was below the detection limit, the numbers were semi-quantified by most probable number (MPN) method for 3 tube assay as described by Oblinger and Koburger (49). Samples were diluted 6-fold in buffered peptone water (BPW) and incubated overnight at 37 °C, and positive samples confirmed by plating triplicate 100 μl samples on MAC agar and incubating overnight at 37 °C.

## Acknowledgments

NJH and SM were supported by a FSA grant (FS101056); BM was supported by a PhD award to NJH, NS, FB and KF; NJH was partly funded by the Rural & Environment Science & Analytical Services Division of the Scottish Government. We are grateful to Susan Verrall and Raymond Campbell (Hutton institute) for assistance with GC-MS and HPLC; David Gally (University of Edinburgh) for use of CL3 facilities.

---

**Supplemental Figure 1** Manual correction of growth rate misfits in DMFIT.

Example of a correction with *E. coli* isolate JHI5039 grown in lettuce leaf lysate, 18 °C. **A)** DMFIT could not fit a non-linear curve on data (n = 193) with a decrease in the stationary phase (R^2^_adj_ = 0.001). **B)** Data was cut off manually (n = 49) to achieve better fits (R^2^_adj_ = 0.996). A complete list of fits including data points are in Supplemental Table 3.

**Supplementa1 Figure 2** Simplified polar metabolic pathways in plants

Interaction between major polar pathways (colour coded) in green leafy plants. Metabolism of carbohydrates degradation (green) is linked to amino acid degradation (dark blue and purple), which feed into the TCA cycle (red). The arrows pointing outside are entries into the non-polar fatty acid pathway. The glutamate group (orange) leads into the urea cycle. The light blue cycle described the acyl chain synthesis. Modified from the metabolomic pathway in *Solanum*, based on Dobson, et al. (14).

## References

1. Baumler, D. J., R. G. Peplinski, J. L. Reed, J. D. Glasner, and N. T. Perna. 2011. The evolution of metabolic networks of E. coli. BMC Syst. Biol. 5:21.

2. Bertani, G. 2004. Lysogeny at Mid-Twentieth Century: P1, P2, and Other Experimental Systems. J Bacteriol 186:595–600.

3. Bertani, G. 1951. Studies on lysogenesis. The mode of phage liberation by lysogenic Escherichia coli. J. Bacteriol. 62:293–300.

4. Bewley, J. D., and M. Black. 1978. Physiology and Biochemistry of Seeds in Relation to Germination: 1 Development, Germination, and Growth. Springer-Verlag Berlin Heidelberg.

5. Buchanan, R. L., and L. A. Klawitter. 1992. The effect of incubation temperature, initial pH, and sodium chloride on the growth kinetics of Escherichia coli O157:H7. Food Microbiol 9:185–196.

6. Buchholz, U., H. Bernard, D. Werber, M. M. Bohmer, C. Remschmidt, H. Wilking, Y. Delere, M. an der Heiden, C. Adlhoch, J. Dreesman, J. Ehlers, S. Ethelberg, M. Faber, C. Frank, G. Fricke, M. Greiner, M. Hohle, S. Ivarsson, U. Jark, M. Kirchner, J. Koch, G. Krause, P. Luber, B. Rosner, K. Stark, and M. Kuhne. 2011. German outbreak of Escherichia coli O104:H4 associated with sprouts. N Engl J Med 365:1763–70.

7. Carter, M. Q., J. W. Louie, D. Feng, W. Zhong, and M. T. Brandl. 2016. Curli fimbriae are conditionally required in Escherichia coli O157:H7 for initial attachment and biofilm formation. Food Microbiol 57:81–89.

8. Charkowski, A. O., J. D. Barak, C. Z. Sarreal, and R. E. Mandrell. 2002. Differences in growth of Salmonella enterica and Escherichia coli O157 : H7 on alfalfa sprouts. Appl Environ Microbiol 68:3114–3120.

9. Crozier, L., P. Hedley, J. Morris, C. Wagstaff, S. C. Andrews, I. Toth, R. W. Jackson, and N. Holden. 2016. Whole-transcriptome analysis of verocytotoxigenic Escherichia coli O157:H7 (Sakai) suggests plant-species-specific metabolic responses on exposure to spinach and lettuce extracts. Front Microbiol 7:1088.

10. Dahan, S., S. Knutton, R. K. Shaw, V. F. Crepin, G. Dougan, and G. Frankel. 2004. Transcriptome of enterohemorrhagic Escherichia coli O157 adhering to eukaryotic plasma membranes. Infect Immun 72:5452–9.

11. Danhorn, T., and C. Fuqua. 2007. Biofilm formation by plant-associated bacteria. Annu Rev Microbiol 61:401–22.

12. Danyluk, M. D., and D. W. Schaffner. 2011. Quantitative assessment of the microbial risk of leafy greens from farm to consumption: preliminary framework, data, and risk estimates. J Food Prot 74:700–708.

13. Deering, A. J., L. J. Mauer, and R. E. Pruitt. 2012. Internalization of E. coli O157:H7 and Salmonella spp. in plants: A review. Food Res Int 45:567–575.

14. Dobson, G., T. Shepherd, R. Marshall, S. R. Verrall, S. Conner, D. W. Griffiths, J. W. McNicol, D. Stewart, and H. V. Davies. 2007. Dordrecht.

15. Elhadidy, M., and A. Álvarez-Ordóñez. 2016. Diversity of survival patterns among Escherichia coli O157:H7 genotypes subjected to food-related stress conditions. Front Microbiol 7.

16. Erickson, M. C., J. Liao, A. S. Payton, C. C. Webb, L. Ma, G. D. Zhang, I. Flitcroft, M. P. Doyle, and L. R. Beuchat. 2013. Fate of Escherichia coli O157:H7 and Salmonella in soil and lettuce roots as affected by potential home gardening practices. J Sci Food Agri 93:3841–3849.

17. Erickson, M. C., C. C. Webb, L. E. Davey, A. S. Payton, I. D. Flitcroft, and M. P. Doyle.2014. Biotic and abiotic variables affecting internalization and fate of Escherichia coli O157:H7 isolates in leafy green roots. J Food Prot 77:872–9.

18. Erickson, M. C., C. C. Webb, J. C. Diaz-Perez, L. E. Davey, A. S. Payton, I. D. Flitcroft, S. C. Phatak, and M. P. Doyle. 2013. Internalization of Escherichia coli O157:H7 following spraying of cut shoots when leafy greens are regrown for a second crop. J Food Prot 76:2052–2056.

19. Erickson, M. C., C. C. Webb, J. C. Diaz-Perez, S. C. Phatak, J. J. Silvoy, L. Davey, A. S. Payton, J. Liao, L. Ma, and M. P. Doyle. 2010. Infrequent internalization of Escherichia coli O157:H7 into field-grown leafy greens. J Food Prot 73:500–506.

20. Erlacher, A., M. Cardinale, M. Grube, and G. Berg. 2015. Biotic stress shifted structure and abundance of Enterobacteriaceae in the lettuce microbiome. PLoS One 10.

21. Franz, E., S. O. Tromp, H. Rijgersberg, and H. J. van der Fels-Klerx. 2010. Quantitative microbial risk assessment for Escherichia coli O157:H7, Salmonella, and Listeria monocytogenes in leafy green vegetables consumed at salad bars. J Food Prot 73:274–285.

22. George, S. M., A. Métris, and J. Baranyi. 2015. Integrated kinetic and probabilistic modeling of the growth potential of bacterial populations. Appl Environ Microbiol 81:3228–3234.

23. Gibson, A. M., N. Bratchell, and T. A. Roberts. 1988. Predicting microbial growth: growth responses of salmonellae in a laboratory medium as affected by pH, sodium chloride and storage temperature. Int J Food Microbiol 6:155–78.

24. Hayashi, K., N. Morooka, Y. Yamamoto, K. Fujita, K. Isono, S. Choi, E. Ohtsubo, T. Baba, B. L. Wanner, H. Mori, and T. Horiuchi. 2006. Highly accurate genome sequences of Escherichia coli K-12 strains MG1655 and W3110. Molecular Systems Biology 2:2006.0007.

25. Holden, N., R. W. Jackson, and A. Schikora. 2015. Plants as alternative hosts for human and animal pathogens. Front Microbiol 6.

26. Holden, N., L. Pritchard, and I. Toth. 2009. Colonization outwith the colon: plants as an alternative environmental reservoir for human pathogenic enterobacteria. FEMS Microbiol Rev 33:689–703.

27. Holden, N. J., F. Wright, K. MacKenzie, J. Marshall, S. Mitchell, A. Mahajan, R. Wheatley, and T. J. Daniell. 2013. Prevalence and diversity of Escherichia coli isolated from a barley trial supplemented with bulky organic soil amendments: green compost and bovine slurry. Lett Appl Microbiol 58:205–212.

28. Hou, Z., R. C. Fink, C. Radtke, M. J. Sadowsky, and F. Diez-Gonzalez. 2013. Incidence of naturally internalized bacteria in lettuce leaves. Int J Food Microbiol 162:260–265.

29. Huang, L. 2012. Mathematical modeling and numerical analysis of the growth of non-O157 Shiga toxin-producing Escherichia coli in spinach leaves. Int J Food Microbiol 160:32–41.

30. Jay, M. T., M. B. Cooley, D. Carychao, G. W. Wiscomb, R. A. Sweitzer, L. Crawford-Miksza, J. A. Farrar, D. K. Lau, J. O’Connell, A. Millington, R. V. Asmundson, E. R. Atwill, and R. E. Mandrell. 2007. Escherichia coli O157:H7 in feral swine near spinach fields and cattle, central California coast. Emerg Infect Dis 13:1908–11.

31. Jensen, D. A., L. M. Friedrich, L. J. Harris, M. D. Danyluk, and D. W. Schaffner. 2015. Cross contamination of Escherichia coli O157:H7 between lettuce and wash water during home-scale washing. Food Microbiol 46:428–33.

32. Jensen, D. A., L. M. Friedrich, L. J. Harris, M. D. Danyluk, and D. W. Schaffner. 2013. Quantifying transfer rates of Salmonella and Escherichia coli O157:H7 between fresh-cut produce and common kitchen surfaces. J Food Prot 76:1530–8.

33. Klerks, M. M., E. Franz, M. van Gent-Pelzer, C. Zijlstra, and A. H. van Bruggen. 2007. Differential interaction of Salmonella enterica serovars with lettuce cultivars and plant-microbe factors influencing the colonization efficiency. ISME J 1:620–31.

34. Koseki, S., and S. Isobe. 2005. Prediction of pathogen growth on iceberg lettuce under real temperature history during distribution from farm to table. Int J Food Microbiol 104:239–248.

35. Koukkidis, G., R. Haigh, N. Allcock, S. Jordan, and P. Freestone. 2017. Salad leaf juices enhance Salmonella growth, colonization of fresh produce, and virulence. Appl Environ Microbiol 83.

36. Leveau, J. H. J., and S. E. Lindow. 2001. Appetite of an epiphyte: Quantitative monitoring of bacterial sugar consumption in the phyllosphere. Proc Natl Acad Sci USA 98:3446–3453.

37. Li, B.-H., X.-F. Ma, X.-D. Wu, and W.-X. Tian. 2006. Inhibitory activity of chlorogenic acid on enzymes involved in the fatty acid synthesis in animals and bacteria. IUBMB life 58:39–46.

38. Linden, I. V. d., B. Cottyn, M. Uyttendaele, G. Vlaemynck, M. Heyndrickx, M. Maes, and N. Holden. 2016. Microarray-based screening of differentially expressed genes of E. coli O157:H7 Sakai during preharvest survival on butterhead lettuce. Agriculture 6:6.

39. Lohaus, G., K. Pennewiss, B. Sattelmacher, M. Hussmann, and K. Hermann Muehling. 2001. Is the infiltration-centrifugation technique appropriate for the isolation of apoplastic fluid? A critical evaluation with different plant species. Physiologia Plantarum 111:457–465.

40. Marshall, J., Y. Rossez, G. Mainda, D. L. Gally, T. Daniell, and N. Holden. 2016. Alternate thermoregulation and functional binding of Escherichia coli Type 1 fimbriae in environmental and animal isolates. FEMS Microbiol Lett DOI: 10.1093/femsle/fnw251.

41. Mayer, C., and W. Boos. 2005. Hexose/pentose and hexitol/pentitol metabolism. EcoSal Plus.

42. McKellar, R. C., and P. Delaquis. 2011. Development of a dynamic growth-death model for Escherichia coli O157:H7 in minimally processed leafy green vegetables. Int. J. Food Microbiol. 151:7–14.

43. McKellar, R. C., and X. Lu. 2004. Modeling microbial responses in food. CRC Press LLC, Florida, USA.

44. Melotto, M., S. Panchal, and D. Roy. 2014. Plant innate immunity against human bacterial pathogens. Front Microbiol 5:411.

45. Merritt, J. H., D. E. Kadouri, and G. A. O’Toole. 2005. Growing and analyzing static biofilms, p. 1B.1.1–1B.1.17, Curr Prot Microbiol.

46. Miller, W. G., M. T. Brandl, B. Quiñones, and S. E. Lindow. 2001. Biological sensor for sucrose availability: relative sensitivities of various reporter genes. Appl Environ Microbiol 67:1308–1317.

47. Mou, B. 2008. Evaluation of oxalate concentration in the U.S. spinach germplasm collection. HortScience 43:1690–1693.

48. Neidhardt, F. C., P. L. Bloch, and D. F. Smith. 1974. Culture medium for enterobacteria. J Bacteriol 119:736–47.

49. Oblinger, J. L., and J. A. Koburger. 1975. Understanding and teaching the most probable number technique. J Milk Food Technol 38:540–545.

50. Painter, J. A., R. M. Hoekstra, T. Ayers, R. V. Tauxe, C. R. Braden, F. J. Angulo, and P. M. Griffin. 2013. Attribution of foodborne illnesses, hospitalizations, and deaths to food commodities by using outbreak data, United States, 1998-2008. Emerg Infect Dis 19:407–15.

51. Pang, H., E. Lambertini, R. L. Buchanan, D. W. Schaffner, and A. K. Pradhan. 2017. Quantitative microbial risk assessment for Escherichia coli O157:H7 in fresh-cut lettuce. J Food Prot 80:302–311.

52. Perez-Rodriguez, F., M. J. Saiz-Abajo, R. M. Garcia-Gimeno, A. Moreno, D. Gonzalez, and S.I. Vitas. 2014. Quantitative assessment of the Salmonella distribution on fresh-cut leafy vegetables due to cross-contamination occurred in an industrial process simulated at laboratory scale. Int J Food Microbiol 184:86–91.

53. Perry, N., T. Cheasty, T. Dallman, N. Launders, and G. Willshaw. 2013. Application of multi-locus variable number tandem repeat analysis to monitor Verocytotoxin-producing Escherichia coli O157 phage type 8 in England and Wales: emergence of a profile associated with a national outbreak. J Appl Microbiol 115:1052–1058.

54. Pignocchi, C., and C. H. Foyer. 2003. Apoplastic ascorbate metabolism and its role in the regulation of cell signalling. Curr. Opin. Plant Biol. 6:379–89.

55. Posada-Izquierdo, G., S. Del Rosal, A. Valero, G. Zurera, A. S. Sant’Ana, V. O. Alvarenga, and F. Perez-Rodriguez. 2016. Assessing the growth of Escherichia coli O157:H7 and Salmonella in spinach, lettuce, parsley and chard extracts at different storage temperatures. J Appl Microbiol 120:1701–1710.

56. Posada-Izquierdo, G. D., F. Perez-Rodriguez, F. Lopez-Galvez, A. Allende, M. I. Gil, and G. Zurera. 2014. Modeling growth of Escherichia coli O157:H7 in fresh-cut lettuce treated with neutral electrolyzed water and under modified atmosphere packaging. Int J Food Microbiol 177:1–8.

57. Presser, K. A., D. A. Ratkowsky, and T. Ross. 1997. Modelling the growth rate of Escherichia coli as a function of pH and lactic acid concentration. Appl Environ Microbiol 63:2355–2360.

58. Presser, K. A., T. Ross, and D. A. Ratkowsky. 1998. Modelling the growth limits (growth/no growth interface) of Escherichia coli as a function of temperature, pH, lactic acid concentration, and water activity. Appl Environ Microbiol 64:1773–1779.

59. Quilliam, R. S., A. P. Williams, and D. L. Jones. 2012. Lettuce cultivar mediates both phyllosphere and rhizosphere activity of Escherichia coli O157:H7. PLoS ONE 7:e33842.

60. Ratkowsky, D. A., R. K. Lowry, T. A. McMeekin, A. N. Stokes, and R. E. Chandler. 1983. Model for bacterial culture growth rate throughout the entire biokinetic temperature range. J Bacteriol 154:1222–1226.

61. Ratkowsky, D. A., J. Olley, T. A. McMeekin, and A. Ball. 1982. Relationship between temperature and growth rate of bacterial cultures. J Bacteriol 149:1–5.

62. Record Jr, M. T., E. S. Courtenay, D. S. Cayley, and H. J. Guttman. 1998. Responses of E. coli to osmotic stress: large changes in amounts of cytoplasmic solutes and water. Trends in Biochemical Sciences 23:143–148.

63. Rico, A., and G. M. Preston. 2008. Pseudomonas syringae pv. tomato DC3000 uses constitutive and apoplast-induced nutrient assimilation pathways to catabolize nutrients that are abundant in the tomato apoplast. Mol Plant Microbe Interact 21:269–282.

64. Rossez, Y., A. Holmes, H. Lodberg-Pedersen, L. Birse, J. Marshall, W. G. T. Willats, I. K. Toth, and N. J. Holden. 2014. Escherichia coli common pilus (ECP) targets arabinosyl residues in plant cell walls to mediate adhesion to fresh produce plants. J Biol Chem 289:34349–34365.

65. Rossez, Y., A. Holmes, E. B. Wolfson, D. L. Gally, A. Mahajan, H. L. Pedersen, W. G. T. Willats, I. K. Toth, and N. J. Holden. 2014. Flagella interact with ionic plant lipids to mediate adherence of pathogenic Escherichia coli to fresh produce plants. Environ Microbiol 16:2181–2195.

66. Sattelmacher, B. 2001. The apoplast and its significance for plant mineral nutrition. New Phytol 149:167–192.

67. Seo, S., and K. R. Matthews. 2014. Exposure of Escherichia coli O157:H7 to soil, manure, or water influences its survival on plants and initiation of plant defense response. Food Microbiol 38:87–92.

68. Seo, S., and K. R. Matthews. 2012. Influence of the plant defense response to Escherichia coli O157:H7 cell surface structures on survival of that enteric pathogen on plant surfaces. Appl Environ Microbiol 78:5882–5889.

69. Shepherd, L. V., J. W. McNicol, R. Razzo, M. A. Taylor, and H. V. Davies. 2006. Assessing the potential for unintended effects in genetically modified potatoes perturbed in metabolic and developmental processes. Targeted analysis of key nutrients and anti-nutrients. Transgenic research 15:409–25.

70. Vital, M., D. Stucki, T. Egli, and F. Hammes. 2010. Evaluating the growth potential of pathogenic bacteria in water. Appl Environ Microbiol 76:6477–6484.

71. Wallace, R. J. 2004. Antimicrobial properties of plant secondary metabolites. The Proceedings of the Nutrition Society 63:621–9.

72. Wright, K. M., S. Chapman, K. McGeachy, S. Humphris, E. Campbell, I. K. Toth, and N. J. Holden. 2013. The endophytic lifestyle of Escherichia coli O157:H7: quantification and internal localization in roots. Phytopathol 103:333–340.

73. Wright, K. M., L. Crozier, J. Marshall, B. Merget, A. Holmes, and N. J. Holden. 2017. Differences in internalization and growth of Escherichia coli O157:H7 within the apoplast of edible plants, spinach and lettuce, compared with the model species Nicotiana benthamiana. Microb Biotechnol 10:555–569

74. Wright, K. M., and N. J. Holden. 2018. Quantification and colonisation dynamics of Escherichia coli O157:H7 inoculation of microgreens species and plant growth substrates. Int J Food Microbiol 273:1–10.

75. Zheng, Y., J. Liu, M. L. Cao, J. M. Deng, and J. Kou. 2016. Extrication process of chlorogenic acid in Crofton weed and antibacterial mechanism of chlorogenic acid on Escherichia coli. J.Environ.Biol. 37:1049–1055.

